# Tidyplots empowers life scientists with easy code-based data visualization

**DOI:** 10.1101/2024.11.08.621836

**Authors:** Jan Broder Engler

## Abstract

Code-based data visualization is a crucial tool for understanding and communicating experimental findings while ensuring scalability and reproducibility. However, complex programming interfaces pose a significant barrier for life scientists. To address this challenge, tidyplots provides a user-friendly code-based interface for creating customizable and insightful plots. With its consistent and intuitive syntax, tidyplots empowers researchers to leverage automated data visualization pipelines while minimizing required programming skills.

Tidyplots is available on CRAN at https://CRAN.R-project.org/package=tidyplots and GitHub at https://github.com/jbengler/tidyplots. The full documentation is available at https://tidyplots.org.

## Main

Data visualization is a crucial component of the data analysis workflow, facilitating efficient data exploration and the extraction of experimental findings, including their direction, magnitude, and robustness. Additionally, it is a key tool for communicating findings in scientific outlets and engaging the community in discussions to validate, corroborate, or challenge results (*1, 2*).

Advances in life sciences methodology have led to a surge in both the volume and complexity of experimental data. As a result, traditional data analysis workflows—often reliant on copy-pasting and manual spreadsheet manipulations—are struggling to meet the demands for higher throughput and the rising standards of reproducibility and transparency. This has prompted the life science community to adopt programmatic data analysis ecosystems. The most widely used tools include R-based packages like the tidyverse (*3*) and ggplot2 (*4*), and Python-based libraries such as Pandas, NumPy, Matplotlib, and Seaborn.

However, despite their utility and power, each of these tools uses a specialized syntax and requires significant coding experience, posing a barrier to adoption by life scientists. Something as simple as a scatter plot with error bars and a statistical test typically requires a considerable amount of code and intricate knowledge of the plotting tool’s internal workings. To address this challenge, this paper introduces the open-source R package tidyplots that was specifically designed to empower life scientists to benefit from automated data visualization pipelines. As such, tidyplots addresses similar needs as ggstatsplot (*5*) and ggpubr (*6*), however instead of extending the ggplot2 syntax, tidyplots introduces a novel interface based on a consistent and intuitive grammar that minimizes the need of programming experience.

The tidyplots workflow is composed of a series of function calls that are connected in one pipeline (**Fig. 1a**). After starting the plot using the *tidyplot()* function, there are three main verbs to construct and modify the plot, namely *add, remove* and *adjust*. Optionally, the user can choose a different *theme, split* the plot into a multiplot layout, or *save* it to file (**Fig. 1a**). As an example, we will use the *study* dataset from the tidyplots package that includes a treatment and placebo group at two different treatment doses and a score measuring treatment success (**Fig. 1b**). Within the *tidyplot()* function we define which variable to use for the x-axis, the y-axis and the color. In the next line, we add the mean value of each group represented as a bar using the function *add_mean_bar()*. This exemplifies a general scheme, in which function names start with an action verb, such as *add*, followed by the statistical entity, such as *mean*, followed by the graphical representation of the entity, such as *bar*. In a similar way, we can add the standard error of the mean as errorbar with *add_sem_errorbar()*, the raw data values as points with *add_data_points()*, and a statistical test as *p* values with *add_test_pvalue()* (**Fig. 1b**). tidyplots comes with over 50 *add* functions that cover the plotting of raw data values, summary statistics, dispersion, distribution, proportion, annotation and statistical comparison (**Fig. 1c,d**). Fortunately, the user does not have to memorize all these names, because auto-completion in the code editor typically suggests fitting functions after typing a few characters. Thus, typing “add_” gives a list of all *add* functions, typing “add_mean” gives a list of all available graphical representations of mean, and typing “add_bar” gives a list of all available statistical entities that can be represented as bars.

**Fig. 1.**
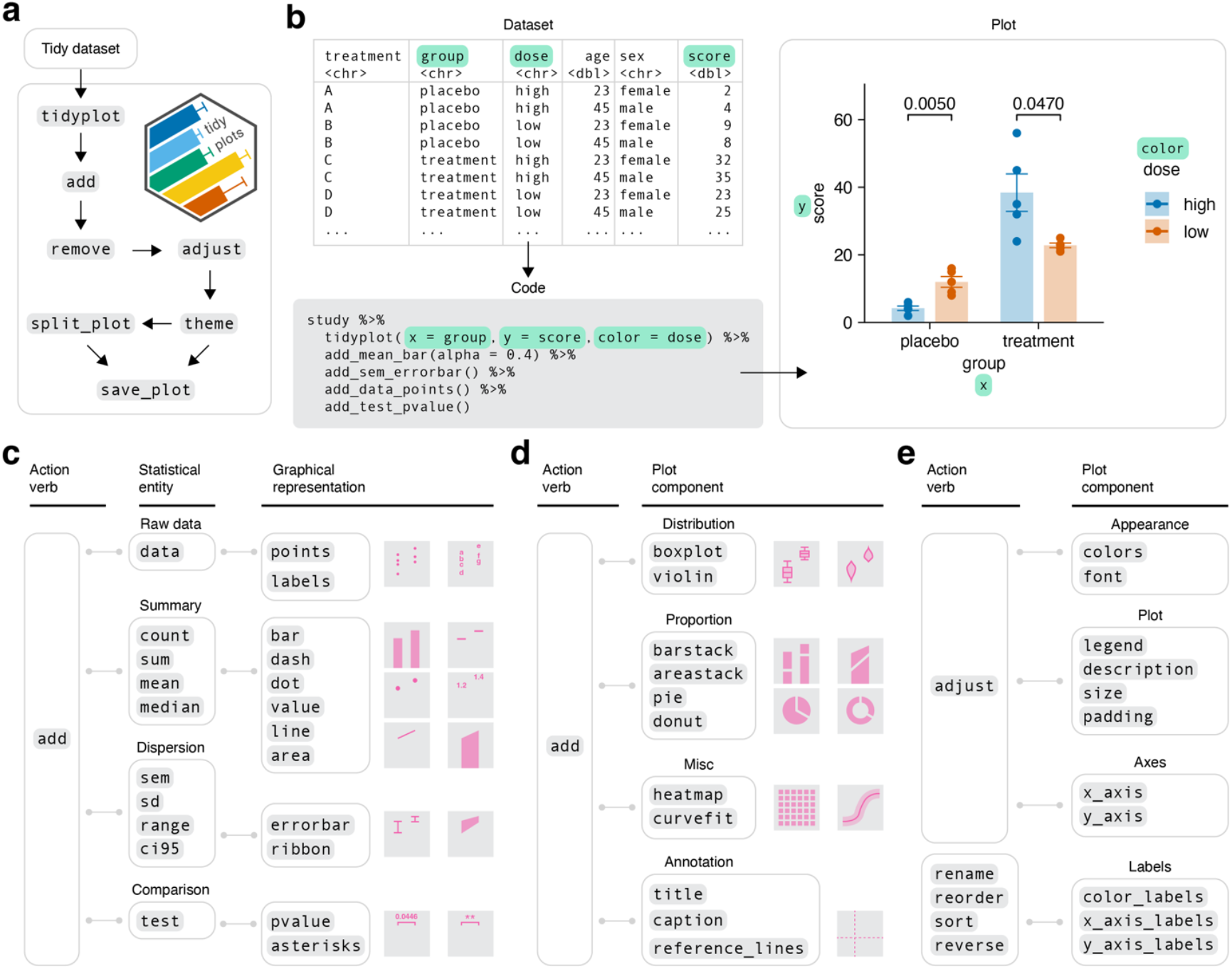
Design principles of tidyplots. **a**, Overview of the tidyplots workflow. **b**, Scheme and example code illustrating the way from the dataset to the plot. **c**, Grammar of *add* functions showing available combinations of statistical entities and graphical representations. **d**, Grammar of *add* functions for predefined plot components. **e**, Grammar of functions to *adjust* the plot.

In the next step, the user can *remove* unwanted plot elements and *adjust* the appearance of the plot, including colors, fonts, legend, titles, size, padding and axes (**Fig. 1e**). A specialized group of functions deals with the color and axis labels that are derived from the dataset itself. Starting with *rename, reorder, sort* and *reverse*, these functions allow to conveniently modify the order and naming of data labels along the axes and the color legend (**Fig. 1e**).

Given its modular architecture, tidyplots can be used to create a wide range of scientific plots (**Fig. 2a**), while its focus on human code readability makes the underlying source code easy to read and write. To illustrate this, we will compare tidyplots to the most popular R plotting package ggplot2. Due to its more granular interface the ggplot2 code is less accessible and requires a deeper understanding of several internal ggplot2 concepts (**Fig. 2b**). In contrast, tidyplots requires considerably less words, characters, function calls and function arguments to achieve plots equivalent to **Fig 2a**, demonstrating a significant reduction of code complexity (**Fig. 2c**).

**Fig. 2.**
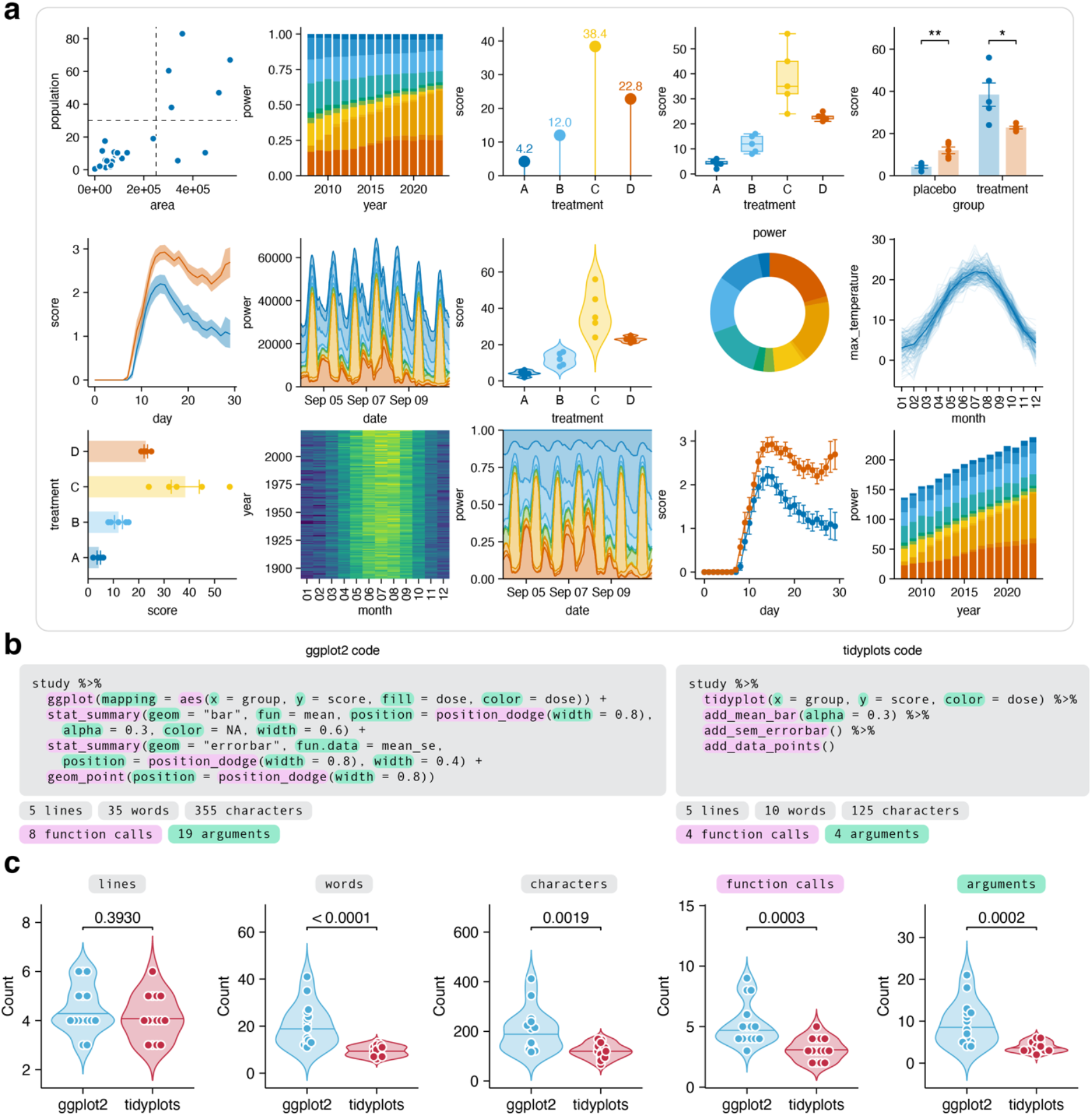
Performance of tidyplots. **a**, Plot gallery. **b**, ggplot2 and tidyplots example code to generate equivalent plots. **c**, Comparison of ggplot2 and tidyplots code across plots in **a**. Statistical analysis was performed by Mann–Whitney *U* test.

Overall, tidyplots provides a powerful and user-friendly solution for creating code-based scientific plots. Beyond its core functionality, tidyplots offers several valuable features, including demo datasets that are ideal for learning and teaching, color schemes optimized for individuals with color vision deficiencies, and thorough documentation with a beginner-friendly getting started guide. Tidyplots also enhances the productivity of experienced users by streamlining the process of creating complex visualizations, allowing them to focus more on data interpretation and analysis. By promoting the use of code-based plotting in the life sciences, tidyplots has the potential to accelerate scientific discoveries and enhance reproducibility and transparency.

## Methods

### Interface design

The interface of tidyplots is heavily inspired by the tidyverse ecosystem (*3*) and its underlying design principles (*7*). Most importantly, tidyplots is designed (i) to maximize human code readability by using natural language elements, (ii) to be consistent across functions and reduce the need to remember special cases, and (iii) to be modular in the sense that a complex task can be broken down into multiple steps. Function names begin with an action verb, such as *add, remove, adjust*, or *save*, clearly indicating the type of action the function performs. For adding summary statistics, statistical entities are given precedence over graphical presentation, encouraging a conscious decision about *what* to plot, such as *mean*, before *how* to represent it, such as *bar*.

Similar to ggplot2 (*4*), which is the plotting framework underlying tidyplots, plots are built by gradually adding layers of information. This modular design enables maximal flexibility and freedom to combine elements in order to compose complex plots. However, a main difference between ggplot2 and tidyplots is that ggplot2 functions represent *nouns* that are added together using the + operator while tidyplots functions represent *verbs* that are combined in a pipeline using the pipe operator. The concept of piping has been widely adopted in R programming because it greatly enhances code readability. By using the pipe, tidyplots aims to increase the consistency across data analysis workflows. Thus, the wrangeling of data sets, the plotting, and the post-processing of the plots, such as multiplot arrangement and saving, can be conveniently combined in one pipeline without the need to switch operators.

Another design decision of tidyplots concerns the prioritization of specialized functions over function arguments. For example, the addition of the mean in several graphical representations could be implemented by one function called *add_mean()* that takes the desired graphical representation, such as *bar, dash*, or *point*, as parameter. Instead, tidyplots implements three individual functions called *add_mean_bar(), add_mean_dash()* and *add_mean_point()*. Thereby, tidyplots encouraged the use of autocompletion in the code editor that gives a list of all available options while typing, thus eliminating the need to consults function documentation. In fact, most tidyplot functions work as intended without any parameters, greatly accelerating the writing of code, improving code readability and facilitating the learning of tidyplots. Following this notion, tidyplots offers over 50 *add* function, that adhere to the same consistent naming scheme.

### Functional programming

The above-mentioned prioritization of specialized functions over function arguments comes with one central challenge in package development, which is code repetition. For example, *add_mean_bar(), add_median_bar()* and *add_sum_bar()* deliver very similar functionality only differing in the statistical summary function they apply. When defining these functions, this would lead to considerable code repetition, which inflates the amount of source code and, more importantly, compromises code maintainability. To address this challenge, tidyplots makes heavy use of function factories (*8*). Function factories are functions that return manufactured functions. For example, the tidyplots function factory *ff_bar()* takes a statistical summary function such as *mean, median*, or *sum*, as input and delivers the manufactured functions *add_mean_bar(), add_median_bar()* and *add_sum_bar()* as output. This approach eliminates code repetition and greatly enhances code maintainability.

### Dependencies

Tidyplots heavily relies on the ecosystem of tidyverse packages (*3*), especially ggplot2 (*4*). However, to deliver specific features, tidyplots also depends on some additional great open-source R packages. These include patchwork (*9*) to enable absolute plot dimensions, ggrastr (*10*) to enable rasterization of layers with too many vector shapes, ggbeeswarm (*11*) to avoid over-plotting by violin-shaped distribution of data points, ggrepel (*12*) to handle overlapping text labels, and ggpubr (*6*) for adding statistical comparisons directly to the plot.

### Benchmarking

For benchmarking code complexity, the code to generate the tidyplots plot gallery (**Fig. 2a**) was analyzed using the tidyverse package stringr and compared to the ggplot2 code needed to generate equivalent plots. Benchmarking metrics included the number of lines, words, characters, functions calls and function arguments required by each plotting tool. The source code for the analysis along with the generated plots are available at https://github.com/jbengler/tidyplots_paper. Statistical analyses of benchmarking metrics were performed using Mann–Whitney *U* test.

## Data availability

All data is part of the tidyplots package, available at https://github.com/jbengler/tidyplots.

## Code availability

Documentation, articles and a getting started guide for life scientists is available at the tidyplots homepage https://tidyplots.org. The tidyplots source code is available at https://github.com/jbengler/tidyplots. The source code for **Fig. 2** is available at https://github.com/jbengler/tidyplots_paper.

## Acknowledgements

I thank Lars Binkle-Ladisch for joint discussions and Manuel A. Friese for his continuous support and feedback. I acknowledge the R and tidyverse communities, whose software and coding paradigms form the foundation of tidyplots. This work was supported by the Gemeinnützige Hertie-Stiftung through a Hertie Network of Excellence in Clinical Neuroscience Fellowship awarded to J.B.E. (grant no. P1200009).

## Author contributions

J.B.E. designed and developed tidyplots, performed the analyses, and wrote the manuscript.

